# Calculating the statistical significance of rare variants causal for Mendelian and complex disorders

**DOI:** 10.1101/103218

**Authors:** Aliz R Rao, Stanley F Nelson

## Abstract

With the expanding use of next-gen sequencing (NGS) to diagnose the thousands of rare Mendelian genetic diseases, it is critical to be able to interpret individual DNA variation. We developed a general method to better interpret the likelihood that a rare variant is disease causing if observed in a given gene or genic region mapping to a described protein domain, using genome-wide information from a large control sample. We implemented these methods as a web tool and demonstrate application to 19 relevant but diverse next-gen sequencing studies. Additionally, we calculate the statistical significance of findings involving multi-family studies with rare Mendelian disease and studies of large-scale complex disorders such as autism spectrum disorder.

## Background

Whole-exome sequencing has enabled the identification of causal genes responsible for causing hundreds of rare, Mendelian disorders in just a few years; however, there remain hundreds, if not thousands, more to be uncovered. The genetic basis has been determined for 4,803 of the rare diseases [1], whereas the number of disease phenotypes with a known or suspected Mendelian basis lies close to 6,419 based on data in Online Mendelian Inheritance in Man (OMIM) [1]. NGS studies are certain to uncover many disease-phenotype relationships in the near future, but for cases involving rare diseases with limited sample sizes, determining causality between phenotypes and novel genes, and distinguishing true pathogenic variants from rare benign variants remains a challenge. Often disease causality of a given rare variant is only clear when additional affected individuals with similar rare variants in the same gene are identified, which can take years to occur due to the rarity of these disorders. Thus, improvements in determining disease causality or likely pathogenicity would greatly enhance efforts to prioritize genes and gene variants for further molecular analysis, even if only a single affected individual was identified.

Variants identified through broad based NGS technologies are typically classified as pathogenic, likely pathogenic, variant of uncertain significance (VUS) or likely benign according to multiple criteria, largely based on prior knowledge about the specific variant. Novel variants are evaluated individually and placed into discrete categories if they meet complex combinations of criteria, which include thresholds for allele frequency, segregation, number of affected unrelated individuals, and known functional relevance [2, 3]. For example, a variant would be deemed pathogenic if the allele frequency threshold falls below a given threshold and the variant segregates with a disorder in at least two unrelated affected families, or if other criteria are met. In brief, variants are evaluated individually based on variant-specific annotations.

An additional source of information that would aid in variant prioritization would be a gene-specific annotation describing mutational burden in the overall population. To illustrate, consider a gene that has very few functional variants in the general population, and several unrelated patients were found to carry distinct protein-altering, rare missense or potential loss-of-function (LOF) variants in the given gene and within a highly conserved protein domain. Under a model for a rare Mendelian disorder caused by highly penetrant variants, we assume that common variants cannot be considered causal, and rare variants in genes intolerant of mutations are deemed highly suspicious of being causal for disease even if no other information is known about the variants. Therefore, knowing the population-wide mutational burden of a given gene for rare variants would be informative.

While there are gene-ranking methods based on other parameters [4], there has been limited work in developing a gene-level ranking systems based on measures for intolerance to mutations in the general population. The Residual Variation Intolerance Score (RVIS) generates a score based on the frequencies of observed common coding variants compared to the total number of observed variants in the same gene or protein domain [5, 6]. Another ranking system, in addition to these parameters, also incorporates the frequency at which genes are found to be affected by rare, likely functional variants, and their findings suggest that disease associations to genes which frequently contain variants should be evaluated with extra caution [7]. Finally, the Exome Aggregation Consortium (ExAC) dataset provides Z scores that describe the degree to which a gene is depleted of missense and LOF variants compared to expected values based on the frequency of synonymous variants [8]. Although these methods may be useful in prioritizing variants in order to highlight those in genes that frequently contain variants, neither results in a score that is directly interpretable in order to calculate statistics about NGS findings and determine the significance of seeing a variant in a given number of affected individuals.

One tool that calculates a *P-*value of finding a true association through clinical exome sequencing, RD-Match [9], allows researchers to calculate the probability of finding phenotypically similar individuals who share variants in a gene through systems such as Matchmaker Exchange. The tool incorporates the probability of an individual having a rare, nonsynonymous variant in a gene by taking the sum of the allele frequencies of all rare (MAF < 0.1%) nonsynonymous variants annotated in ExAC [8]. With higher MAF thresholds and large population sizes, this is problematic because an individual may have multiple variants in a gene that frequently contains rare variation, causing one to overestimate the fraction of the population carrying rare variants in the gene, hence the fixed, low MAF threshold. Furthermore, this tool is applicable to studies in which the affected individuals are selected based on phenotype as well as the prior knowledge that they share rare variants in a given gene. Finally, RD-Match does not allow researchers to customize variant filtering thresholds according to the disease model with regards to minor allele frequency or predicted consequence such as LOF or missense variant.

Here we describe a method, named SORVA for Significance Of Rare VAriants, for ranking genes based on mutational burden. In addition to incorporating information from variant allele frequencies, we use population-derived data to precompute an unbiased, easily interpretable score, which allows one to calculate the significance of observed and novel rare variants and their potential for being causal of disease. One may then answer the question: what is the probability of observing missense variants in three out of ten unrelated affected individuals, for example, given that only one in a thousand individuals in the general population carry a missense variant in the gene? Essentially, a model can be constructed to estimate the probability of drawing *n* unrelated families with similar biallelic genotypes by chance from the general population [10]. Conversely, if one has a large list of variants of unknown significance, the significance level may be useful in prioritizing variants within the same category of pathogenicity, and in improving the interpretation of variants in studies of Mendelian genetic disorders. This approach is useful for single individuals and small family units, but of course power improves with larger numbers of affected individuals.

## Results

For calculating the significance of seeing variants within a gene when sequencing multiple individuals affected for a rare, presumably Mendelian disorder, we first calculated the frequency of observing a variant in each gene in an individual within the population by using a large control dataset and collapsing variants in each gene. Calculations are based on data from 2504 individuals in the 1000 Genomes Project phase 5 dataset, which contains data from individuals from five “superpopulations” (European, African, East Asian, South Asian, and ad-mixed American). We repeated the analysis for variants filtered according to various minor allele frequency and protein consequence thresholds that researchers may use when filtering variants. First, we filtered out common variants that met various minor allele frequency (MAF) thresholds used in the literature and others: 5%, 1%, 0.5%, 0.1% and 0.05%. We then filtered rare variants according to two scenarios before collapsing variants across genes: 1) we included all protein-altering variants, i.e. those that cause a nonsynonymous change in the protein transcript or have a more deleterious consequence, and 2) we filtered for potential loss-of-function (LOF) variants, i.e. splice site, stop codon gain and frameshift variants.

Below, we present general findings in population and molecular genetics that can be gleaned from the dataset, and illustrate how the dataset can be used in multiple studies, as a control group to vet candidate genes and variants.

### Population differences

Of 18,877 genes that are in the union of the Ensembl and RefSeq gene sets, most genes contained heterozygous or homozygous missense variants in individuals in all populations; only 2.3% contain no rare variants (MAF < 5%), and 1.0% of genes have an identified variant in only a single population. Lowering our MAF threshold does not decrease the number of genes much. Although, filtering variants to include only LOF variants reduces the number of genes containing variants in the dataset to 9641, or 51.1% of genes in the dataset. (Figure 1) These results demonstrate that choosing the correct MAF threshold is not nearly as important as identifying the correct protein consequence threshold to use when filtering variants. For instance, including all missense variants when LOF variants are generally causal for a given disease would reduce power to detect the gene associated with the disease.

**Figure 1.**
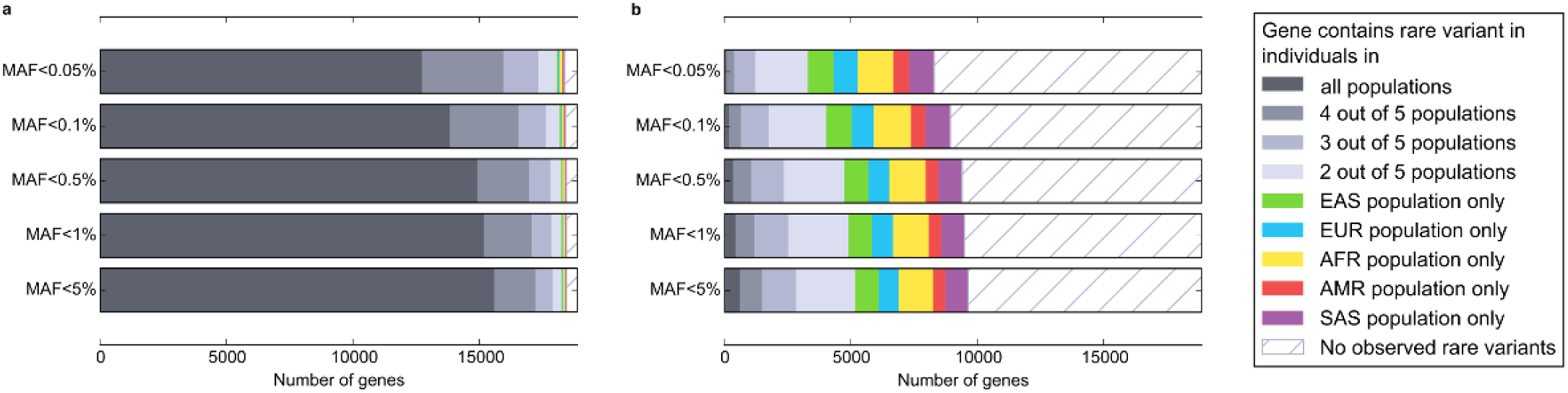
The proportion of genes (n=18877) mutated in individuals in various populations. A gene was considered mutated if at least one individual was heterozygous or homozygous for an uncommon or rare (MAF < 5%) protein-altering (nonsynonymous or potential loss-of-function) variant anywhere in the gene. Abbreviations: EUR, European. AFR, African. EAS, East Asian. SAS, South Asian. AMR, ad-mixed American.

The number of individuals who carried a heterozygous or homozygous variant in a given gene was generally higher in the African population compared to other populations (Figure 2a), which is expected given that African individuals are observed to have up to three times as many low-frequency variants as those of European or East Asian origin [11], which reflects ancestral bottlenecks in non-African populations [12]. Conversely, regarding genes for which the number of individuals with a rare variant in the gene differed between populations, the genes having the greatest difference between populations tended to diverge most in the African population. (Figure 2b) Genes whose mutational burden diverges most between populations are significantly enriched for a large number of biological functional terms, including glycoprotein, olfactory transduction and sensory perception, cell adhesion, various repeats, basement membrane and extracellular matrix part, cadherin, microtubule motor activity, Immunoglobulin and EGF-like domain. It is important to note differences between populations, because, in many cases, researchers would be advised to use control populations similar to their study population. However, if a gene is associated with a severe, childhood-onset disorder in one population, it is likely to be associated with disease in other populations, as well, and knowledge that a gene frequently contains variation in African populations would be useful in prioritizing candidate genes even if one is studying variation in another population. In this case, such information would point towards reduced likelihood for disease association.

**Figure 2.**
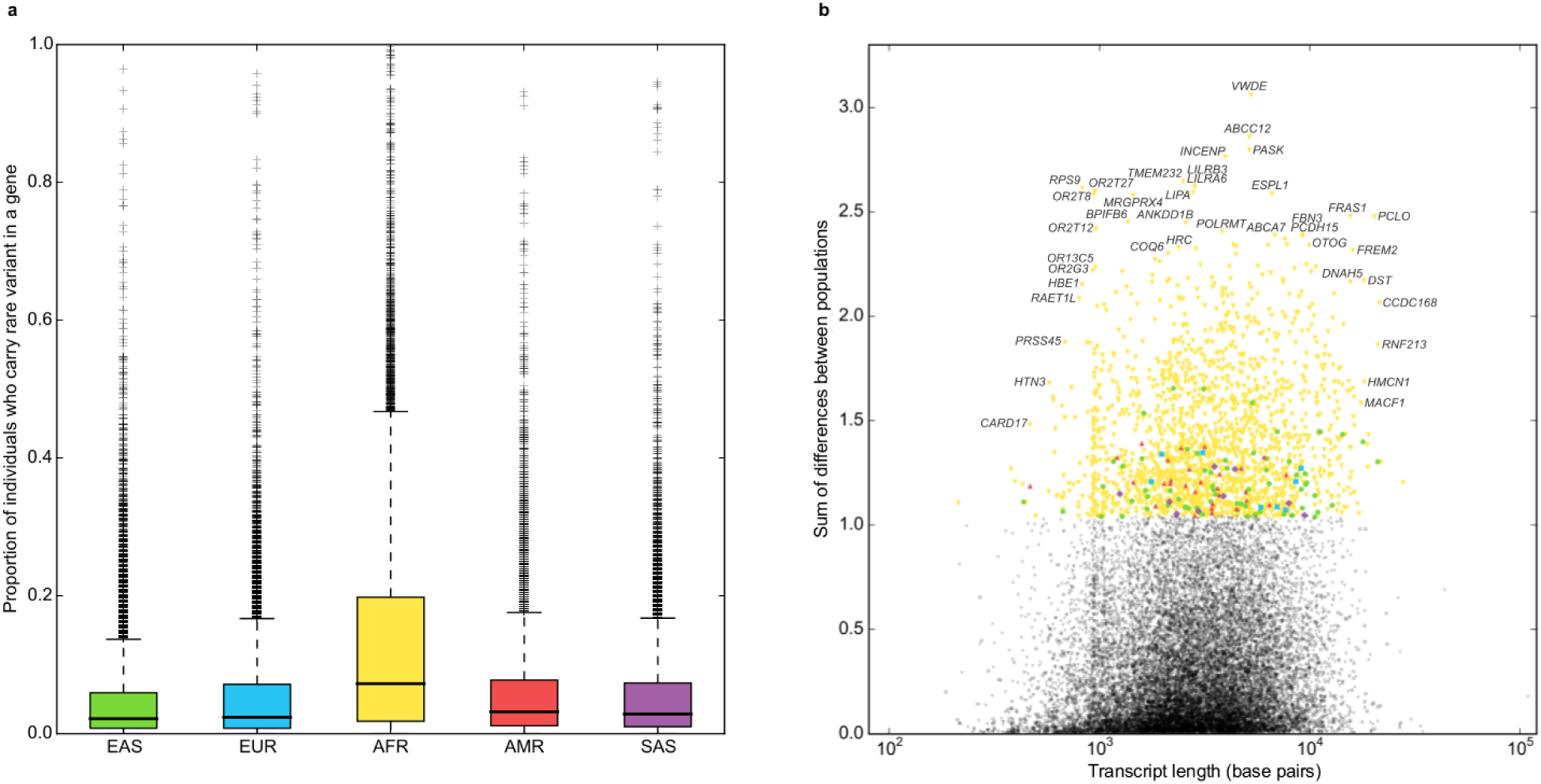
Population differences between the number of individuals mutated for a gene between populations. **(a)** Each data point in the histogram represents the proportion of individuals within a population who are heterozygous or homozygous for an uncommon (MAF < 5%) missense variant in a given gene. **(b)** The number of individuals carrying uncommon variants in a gene differs between populations. We plotted the variance of the count for each gene and colored high-variance genes to denote which population differed most from the mean.

### Properties of known disease genes

To determine whether calculating the frequency of individuals who have a rare variant in a given gene in the general population may be helpful in determining which genes are more likely to cause disease, we compared the counts between multiple categories of genes: a) “essential” genes, defined as genes essential for cell survival in human cell lines, b) genes in which variants are known to cause autosomal dominant disorders, c) genes in which variants are known to cause autosomal recessive disorders, d) genes in which variants are known to cause X-linked disorders, and e) all other genes. As expected, fewer individuals carry rare, protein-altering or LOF variants in genes known to cause Mendelian disorders compared to other genes, and genes associated with X-linked disorders tend to be least tolerant of mutations (Figure 3; Additional file 1). Although frequency counts overlapped between gene categories for every variant filtering threshold, clusters were most differentiated when plotting the proportion of individuals who are heterozygous for rare LOF variants in a gene. Furthermore, the differentiation between clusters increased as the MAF threshold became more stringent, as the datasets became enriched for deleterious variants that can only subsist at a low allele frequency in a population due to selective pressure. (Additional file 1)

**Figure 3.**
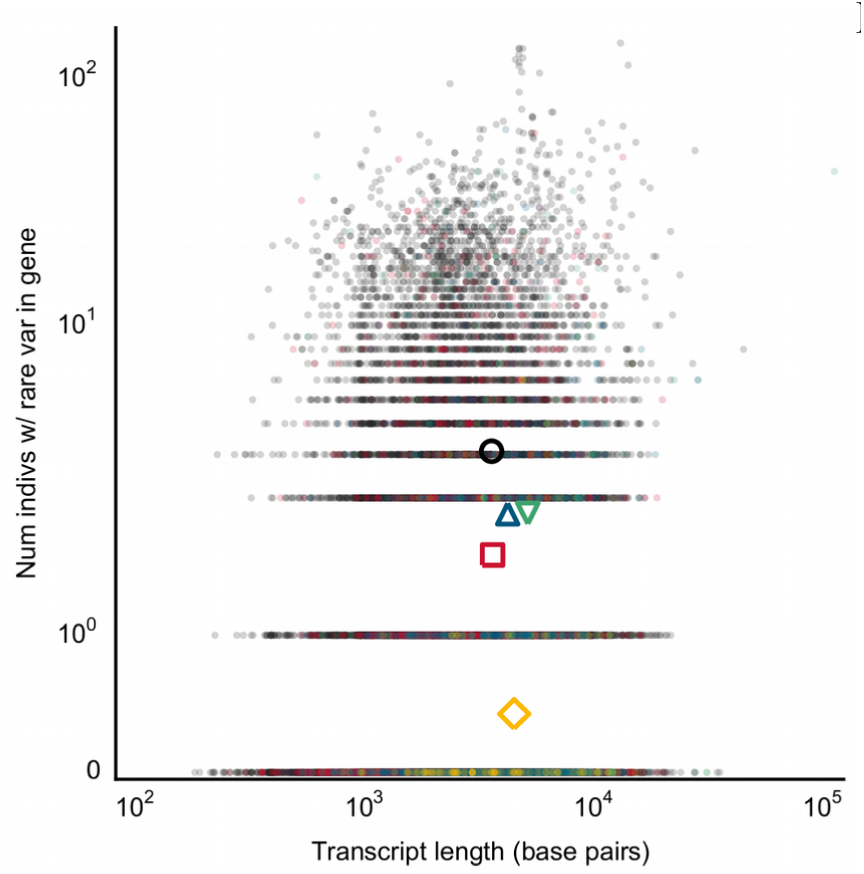
The number of individuals heterozygous for a rare (MAF < 0.5%) potentially LOF mutation in a gene. Each data point represents a single gene, mutated in the aggregate population (n=2504 individuals). Genes are grouped according to whether they are an essential gene, or are known to cause autosomal dominant, autosomal recessive or X-linked disease. Colored shapes indicate the centroids of each group. Abbreviations: nonsyn, nonsynonymous. LOF, loss-of-function. AD, autosomal dominant. AR, autosomal recessive. XL, X-linked.

Previous research suggests that 2.0% of adults of European ancestry and 1.1% of adults of African ancestry can be expected to have actionable highly penetrant pathogenic (including novel expected pathogenic) or likely pathogenic single-nucleotide variants (SNVs) in 112 medically actionable genes [2]. We find that a larger proportion of 1000 Genomes Project individuals—5.8% of European individuals and 3.3% of African individuals—are heterozygous or homozygous for extremely rare (MAF < 0.0005) LOF variants in these 112 genes, highlighting the large number of benign variants that are found in the population at low allele frequencies and should be filtered out by manual curation.

### Depletion of variants in regions mapping to specific protein domains

It has been suggested previously that collapsing variants by protein domain could lead to improved gene-based intolerance scoring systems, as certain regions of the gene could be much more constrained than others [5]. We incorporated data for 322,772 protein domains from Interpro [13] and calculated the average number of individuals who have a variant in any given type of protein domain (Additional file 2), after filtering for rare (MAF < 0.5%), heterozygous LOF variants. Protein domains that are highly constrained, well covered during exome sequencing and rarely contain variants despite their large size include the Family A G protein-coupled receptor-like protein domain (Superfamily: SSF81321), which is found in 660 genes and has a mean length of 965 base pairs; none of the 2,504 individuals carry rare variants in the region mapping to this protein domain. Other highly constrained protein domains that occur throughout the human genome include Glutamic acid-rich region profile (PfScan: PS50313), Proline-rich region profile (PfScan:PS50099), Immunoglobulin (Superfamily: SSF48726), and Cysteine-rich region profile (PfScan: PS50311). (Additional file 2) If an NGS study finds that affected individuals have rare variants in variation intolerant protein domains such as those listed, the variants would become highly suspicious of being causal.

We also calculated whether specific genes contain protein domains that are significantly depleted of variation, given the frequency of variants in the gene overall. Filtering out protein domains in genes with no variants and those with missing information reduced the dataset to 67,138 protein domains in 7,004 genes. The number of rare (MAF < 0.5%), heterozygous LOF variants per individual in a protein domain are significantly lower than expected for 77 protein domains in 26 genes after correcting for multiple testing by the number of genes. (Figure 4) Functional enrichment analysis in DAVID revealed that the most significant biological functions in the gene list were related to tubulin-tyrosine ligase activity (P=0.015), and G-protein coupled receptor, rhodopsin-like superfamily (P=0.05). Depletion values for all protein domains may be found in Additional file 3. Information about whether a protein domain is significantly depleted of variation despite being in a gene with frequently observed variation may be useful in distinguishing between pathogenic and benign rare variants within genes containing regions under different degrees of evolutionary constraint.

**Figure 4.**
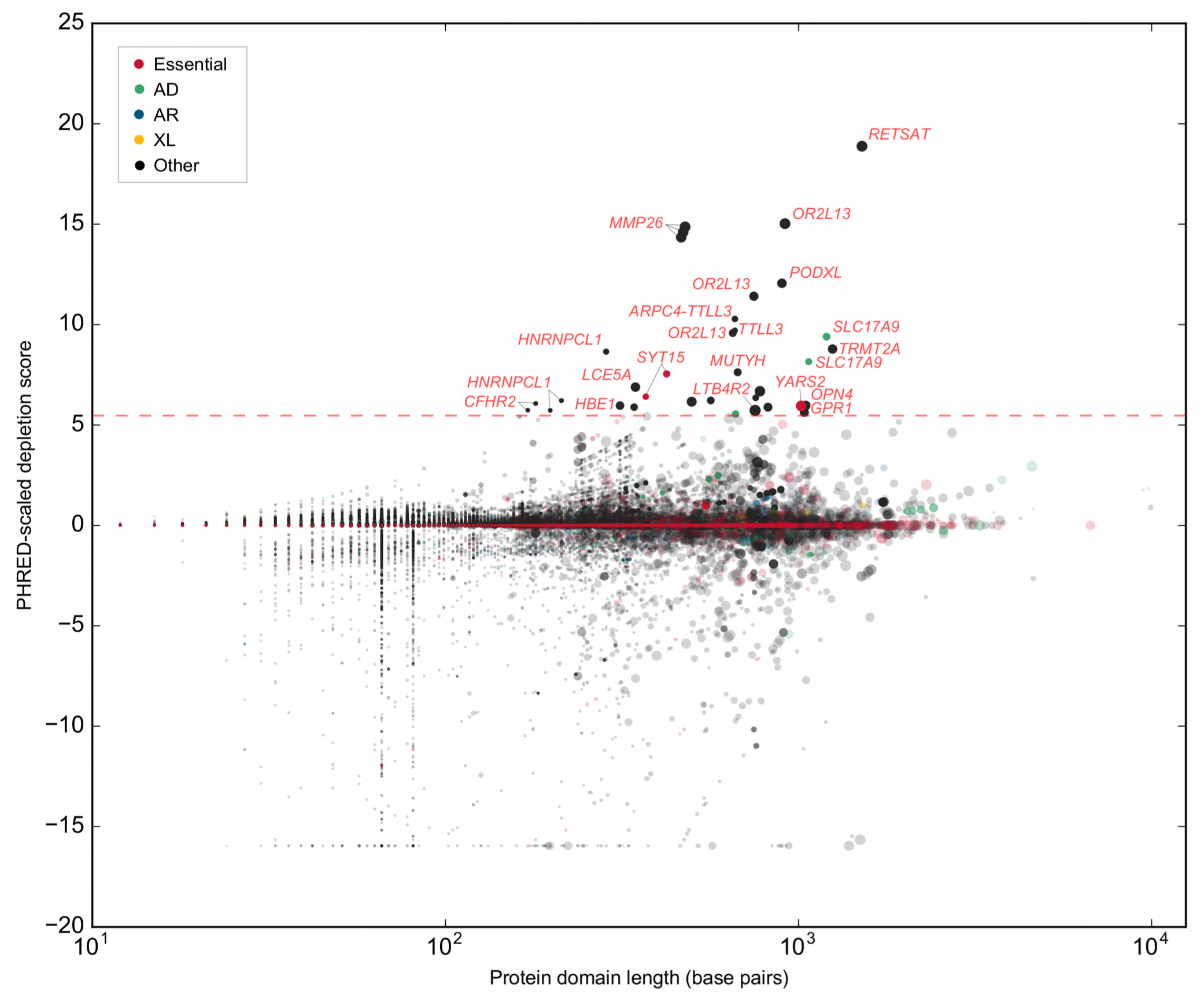
Depletion of rare, heterozygous LOF variants in regions mapping to protein domains. We plotted scaled protein domain depletion scores for each domain mapping within a gene; high scaled scores indicate that a protein domain is depleted of rare (MAF < 0.5%) mutations compared to the rest of the gene. Darkened points above the red dashed line represent protein domains that are significantly depleted of mutations after correcting for the number of genes remaining after filtering. Larger points indicate protein domains with a greater length in proportion to the transcript length. Points are colored if the protein domain is within a gene that is an essential human gene or is causal for a Mendelian disorder. Abbreviations: AD, autosomal dominant. AR, autosomal recessive. XL, X-linked.

### Significance of findings in multi-family studies of rare genetic disorders

Below, we present methods for multiple study designs to calculate the significance of observing a given variant in a given gene. In the simplest case, a study involving a single family, calculating the *P*-value is relatively simple. Consider a case of a severe, pediatric-onset Mendelian disorder, in which both parents and the affected child are sequenced to identify the causal variant. If only *de novo* variants are identified within a putative gene, one can easily estimate the probability of at least one *de novo* mutation occurring in a gene by random chance; one could multiply the per-base mutation rate by the length of the gene transcript and make adjustments to account for CpG content related variation in mutation rates (Additional file 4).

In studies that identify both *de novo* and inherited variants in more complex family structures, calculating the significance of a variant is more complex. First, we generalize the equation for calculating the significance of observing a *de novo* mutation in a gene for studies involving multiple families. The P-value of observing independent *de novo* events in the same gene in *s* out of *n* individuals is

*P*=1−*BinomCDF*(*s*-1, *n*, *l_tx_ dc*)
if multiple families are sequenced. Consider the following example.

Clinical exome sequencing (CES) in four independent families identified *de novo* nonsense mutations in *KAT6A* in all probands displaying significant developmental delay, microcephaly, and dysmorphism [14]. *De novo* nonsense mutations arising in this gene in all four individuals is highly unlikely by chance (*P* = 2.66 × 10^−12^), and the statistical findings would support *KAT6A* as highly suspicious for causing the disorder. Further experiments and the identification of multiple other affected individuals by a separate study [15] confirmed this result.

If inherited variants are also observed in a gene, calculating the statistical significance of findings requires incorporating information about the number of individuals who carry a variant in the particular gene in the general population. The frequencies of the number of individuals who contain rare variants in a given gene or protein domain for various filtering thresholds may be queried through our online database called SORVA (https://sorva.genome.ucla.edu). (Additional file 5) Researchers can select the variant filtering thresholds identical to those used in hard filtering variants in a given study. Minor allele frequency thresholds range from 5%, useful for studies involving more common, complex disorders where less stringent filtering criteria are used, to 0.05% for studies involving extremely rare disorders. Then, knowing the expected number of individuals who carry a variant in the gene or protein domain in question, one can calculate the significance (α < 0.05) of seeing the observed number of singletons (variants observed in a single individual), doubletons (variants observed in two individuals within a single family) or more complex cases as follows.

Let *f_hom_* be the fraction of individuals in the general population with a homozygous variant in a gene or protein domain. Then, the *P*-value of seeing *k* families with a homozygous variant, out of *n* total families with identical relatedness is

*P*_*k,n,r*_=1-*Binom CDF*(*k*−1, *n*, *rf*_*hom*_)

where *r* is the coefficient of relationship[16] or the fraction of the genome shared between affected family members and BinomCDF denotes the binomial cumulative distribution function.

If the affected individuals are heterozygous for the putative variants, the *P*-value is

*P_k,n,r_*=1-*Binom CDF*(*k*-1, *n*, *rf_both_*)

where *f_both_* is the probability of an individual having either a heterozygous or homozygous variant in the gene of interest.

If multiple families and unrelated individuals had been sequenced with different degrees of relatedness, the *P*-value can be obtained by multiplying the probabilities calculated for families grouped by relationship coefficients. To illustrate, consider that we have sequenced 5 singletons (unrelated individuals), three doubletons sharing ½ of their genomes (e.g. siblings concordant for disease status), and a tripleton sharing 1/16 of their genome (e.g. certain distantly related individuals). After variant filtering, we note that the distantly related individuals and four unrelated patients carry rare variants in the same gene, and we calculate the significance of seeing rare damaging variants in four singletons and one tripleton as

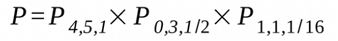

given the family structures of the individuals we had sequenced.

The *a priori* probability *p* can be queried from the SORVA dataset online, and standalone computer software for obtaining *p* and calculating the *P*-value based on the methods described herein is also available on our website.

### Significance of findings in large-scale studies of complex disorders

In complex disorders where most of the genes contributing to risk remain unknown, our dataset may be used to provide additional evidence supporting novel gene findings and provides a simple method to calculate the significance of observing variants in a given gene in a large-scale study. As an example, several large-scale whole-exome sequencing (WES) studies have been carried out to-date in trios and quads to elucidate causal genes underlying autism spectrum disorders (ASD) [17–22]. However, genes identified as containing *de novo* variants rarely overlap between studies, raising the question of how many genes are truly causal and how likely genes are to be identified as associated with autism by chance in these studies as well as others. We assessed the number of individuals carrying rare (MAF<0.1%), heterozygous LOF variants in 1145 genes cumulatively associated with ASD by more than a dozen studies, meta-analyses and reviews [20, 23–41]. There was no significant difference between the distribution of values and that of all genes, and assuming that truly causal genes are more intolerant of rare LOF variants, our findings support the hypothesis that many genes could have been randomly associated with the disorder. (Figure 5, Additional file 6) Furthermore, there are 19 putative autism genes in which >0.5% of individuals carry rare, LOF variants. These genes are likely to be false positives, because no single gene contributes to a large proportion of autism cases. Our results highlight the need to perform statistical validation of findings involving genes associated with complex disorders.

**Figure 5.**
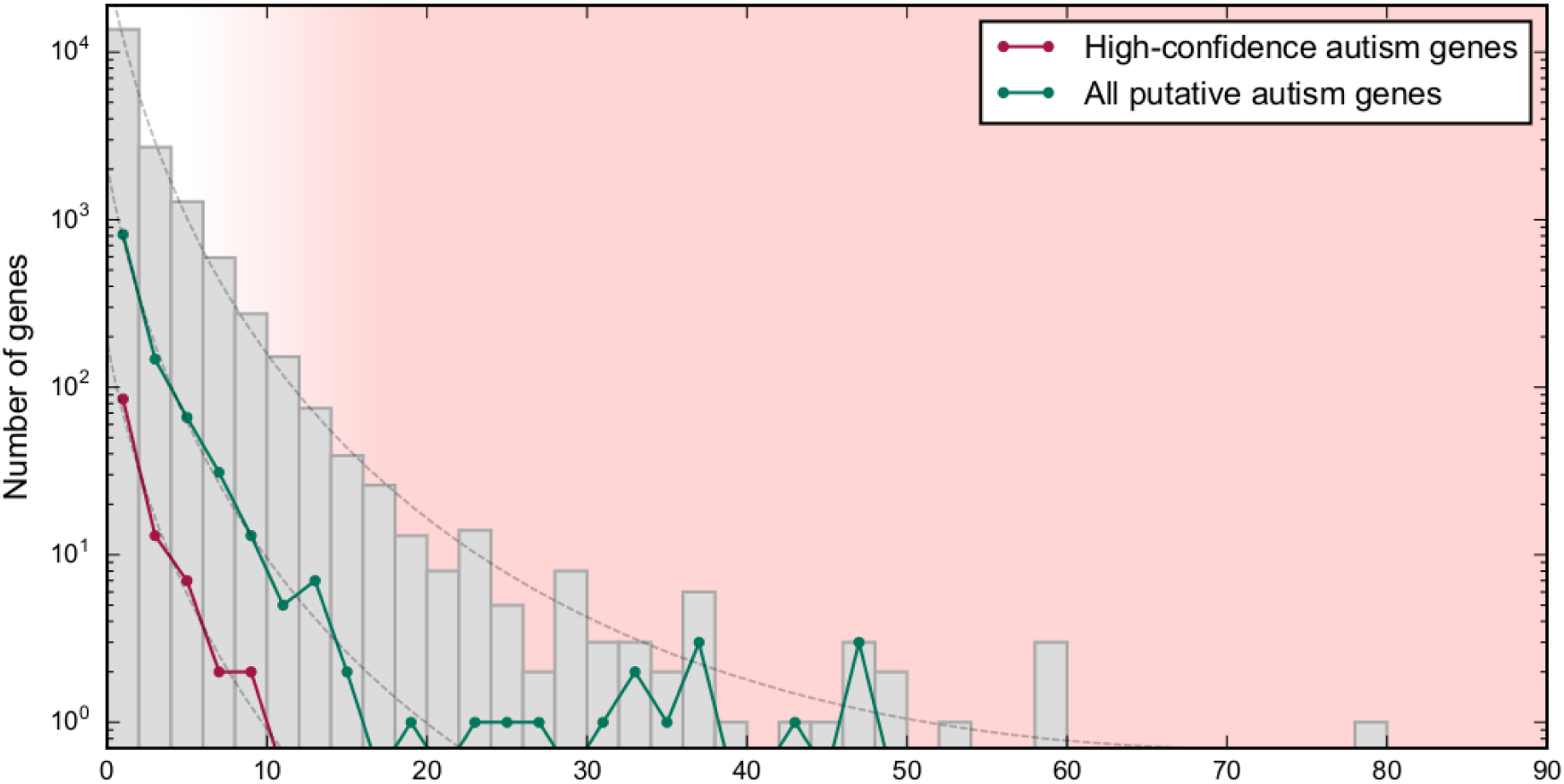
Histogram of the number of individuals with rare LOF variants in putative autism genes. The distribution of the number of individuals with a rare variant (MAF < 0.1%) in all genes is nearly identical to the distribution for putative autism genes (N=1145) and high-confidence autism genes (N=109) (dashed lines), suggesting that the genes may have been associated with autism by chance. Genes that frequently contain rare LOF variants in the population (red shaded region) are unlikely to be causal for ASD.

Appropriately, several WES studies on ASD calculate the significance of their findings. For example, Sanders *et al.* demonstrate in a study which identifies *de novo* coding mutations in 928 individuals that finding two independent *de novo* mutations in a single gene is highly unlikely by chance, and this occurring is viewed as evidence for association between ASD and the gene *SCN2A* (sodium channel, voltage-gated, type II, α subunit) [21]. Neale *et al.* also consider the probability of seeing two independent *de novo* mutations in a single gene when evaluating their findings [18]. Iossifov *et al.* (2012) demonstrates that disrupted genes are significantly enriched for FRMP-associated function; however, they also highlight several individual non-FRMP-associated genes based on their plausibility to cause an ASD phenotype but make no attempt at applying statistics when considering these. In fact, *de novo* mutations in genes may have arisen in these genes by chance and are not causal [17]. To validate our methods, we validated findings by O'Roak *et al.* (2012) [20], who reported *de novo* variants as well as inherited LOF variants in ASD cases. In a targeted sequencing study of 44 candidate genes in 2,446 ASD probands, the authors found that six individual genes (*CHD8*, *GRIN2B*, *DYRK1A*, *PTEN*, *TBR1*, and *TBL1XR1*) had evidence of mutation burden for *de novo* variants, for which they calculated the *P*-value using simulations. Applying our methods, we find that more cases carry *de novo* and inherited variants than expected by chance in 5 out of the 6 genes. (Table 1). The gene *TBL1XR1*, which was borderline significant in the original study, was not significant using our calculations. Furthermore, one additional gene was found to be significant using our method, due to the fact that our method also incorporates information about inherited variants: given that 0.08 % of individuals in the population have a rare (MAF <= 0.05) loss-of-function (LOF) variant in the gene *ADNP*, the significance of seeing *de novo* variants in 2 individuals and additional variants in 1 individual out of a total of 2,446 sequenced singleton individuals is significant after correcting for multiple testing (*P* = 5.15 × 10^−4^). To summarize, our methods approximate *P*-values obtained using more complex and computationally intensive methods such as simulations, with the advantage that it incorporates information about both inherited and *de novo* variation, and the fact that it incorporates precomputed population level data makes our methods easy to apply to calculating the statistical significance of observing rare variants in a given gene.

**Table 1.**
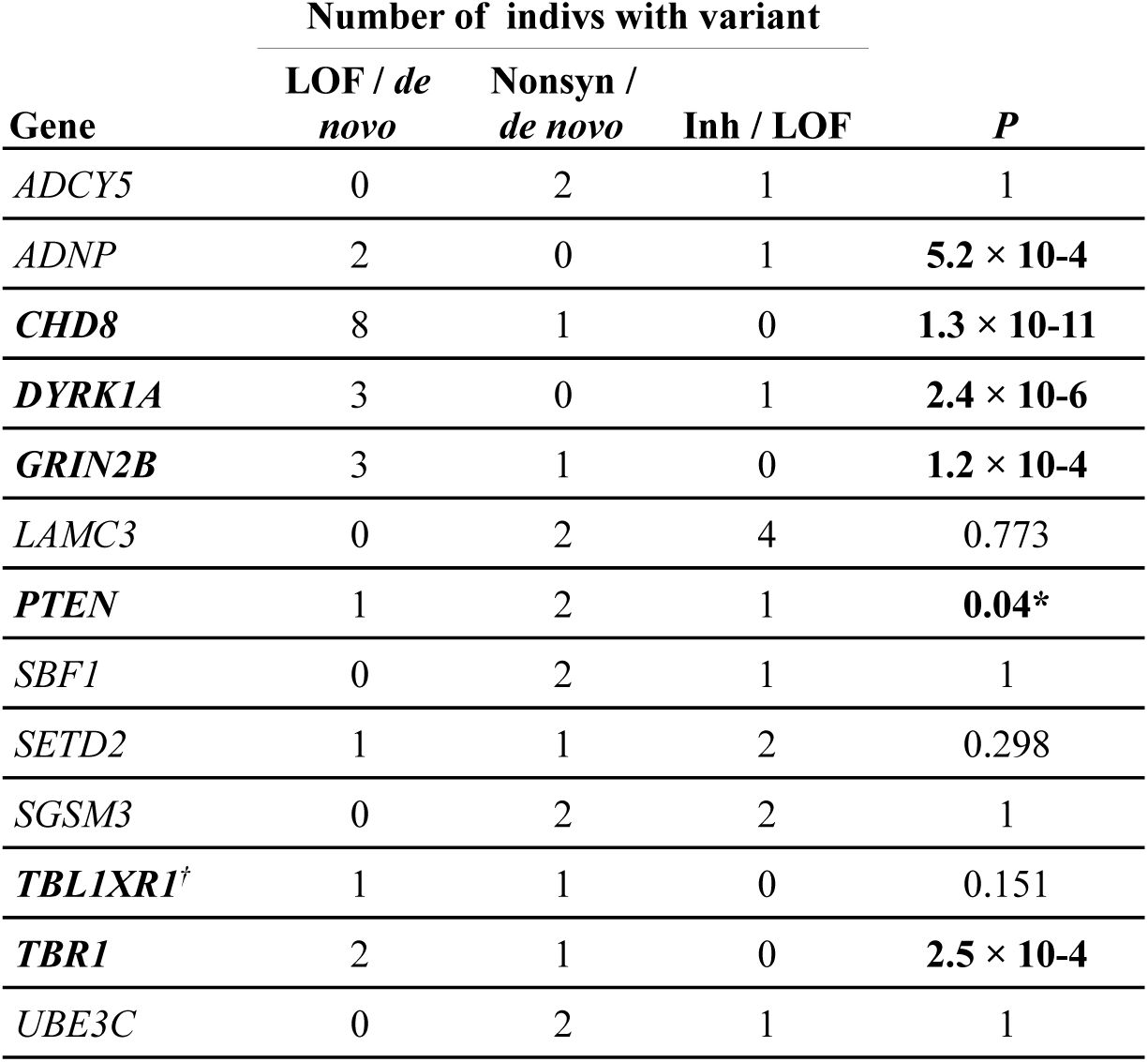
Validation of mutational burden findings in autism genes Number of indivs with variant

In a targeted sequencing study of 44 candidate autism genes in 2446 individuals [20], 12 genes contained both recurring *de novo* variants and inherited LOF variants in multiple individuals, or had evidence of excess mutation burden of *de novo* variants. Gene names that are in bold were statistically significant in the original study. P-values calculated using our methods validate findings by O'Roak *et. al* (2012) for 5 out of 6 genes. \**P*-value was significant if we included nonsynonymous variants. Inherited nonsynonymous variants were not reported in the original study, hence the *P*-value is conservative. †Borderline significant in the original study. Abbreviations: nonsyn, nonsynonymous variant or single amino acid deletion; LOF, loss-of-function variant; Inh, inherited.

### Applications in predictive genomics

If a genetic disease is associated with the presence of variants in a given gene, information about the variants in the gene in affected individuals and in population controls can be used to more accurately assess the probability that a person will develop a disease given their genotype.

Consider a randomly chosen person from the general population who is undergoing prenatal genetic testing. Define *A* as the event that their child will be born with a disease, and *B* as the event that the child carries a rare, LOF variant in a given gene associated with the disease. For many heterogeneic Mendelian disorders, studies of large cohorts provide information regarding the relative contribution of individual causative genes and the genotype–phenotype correlations, giving us the conditional probability *P(B|A)*. The term *P(A)* can be defined as the disease incidence, and the value of *P(B)*, or the proportion of individuals carrying a rare, LOF variant in the gene, can be queried from our dataset. Then, according to Bayes' theorem

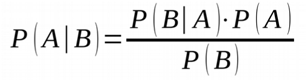

we can calculate that the probability that the child will have the disorder. The following example illustrates such an application.

Consider that prenatal testing identified that a fetus is compound heterozygous for novel variants in the gene *POMGNT1*, which suggests a possible phenotype of congenital muscular dystrophy (CMD). It is known that 53% of patients with CMD have homozygous or compound heterozygous variants in one of six known CMD genes, 10% have homozygous or compound heterozygous variants in *POMGNT1*, and the incidence of CMD is estimated to be 1:21,500 [42, 43]. Since most mutations observed in affected individuals are novel and are not found in healthy population controls, we will assume a low MAF threshold of 0.1% for variant filtering. At this threshold, 2 out of 2504 individuals (0.08 %) in our dataset have a rare protein-altering variant in the gene *POMGNT1*, therefore *P(B)*=0.0008, and we calculate that the positive predictive value (PPV), the probability that the child will have the disease given a positive test result, is roughly 1.0%. Using this method, sensitivity, the probability *P(B|A)*, is quite low (10%); whereas specificity is high (1-P(B) = 99.9%). If we aggregate data for all known CMD genes, we can increase sensitivity to 53% with a negligible decrease in specificity, due to the fact that the other CMD genes contains very few, in any variants in our dataset. This example highlights that sensitivity greatly depends on the proportion of cases that can be explained by variants in a given set of genes. This type of analysis thus has implications for interpretation of broad NGS-based prenatal testing and can be extrapolated as well to preconception testing and risk to potential children.

It is important to note that the extreme numbers involved—the very low prevalence of a disorder and in many cases, the fact that no individual on the 1000 Genomes Project dataset had been observed with variants in a gene, i.e. the lack of previous false-positive results—make it difficult to compute the PPV. A previous study suggests that the latter “zero numerator” problem can be solved using a Bayesian approach that incorporates a prior distribution describing the initial uncertainty about the false-positive rate [44]. Alternatively, the number of rare LOF variants observed in a gene has been published as part of the ExAC dataset, which contains information about 60,706 individuals [8]. Although only nonsense or splice site variants were included in the LOF classification, and they only include values for a single MAF threshold of 0.1%, the number can be used a rough estimate for *f.* Furthermore, if even the ExAC count is zero, we can assume that *f* is less than 1/60706, or 3/60706 if we are being conservative.

To summarize, for monogenic disorders and disorders where there exist detailed phenotype-genotype correlation data, our dataset will provide the denominator in the equation to calculate the probability that an individual with a rare variant in a known disease gene will have a rare genetic disorder. As further research uncovers novel gene-disease associations, and as we increase the size of the public dataset from which *P(B)* values can be calculated, we can update expected false-positive rates and calculating PPVs will become increasingly accurate. As illustrated, our methods will be be useful for applications in predictive genomics, including prenatal testing and testing for late-onset genetic disorders.

### Comparison to other gene ranking methods

We applied our method to calculate the significance of several previous studies' findings [45–72]. In all studies where the Mendelian disorder was found to be caused by inherited disease variants (N=21) [45–62, 70–72], findings were confirmed to be significant using our methods, and in 15 out of 21 studies, *P*-values were highly significant (*P*< 0.0001). (Table 2, Additional file 7) In many studies, initial exome sequencing in a limited number of individuals is followed by sequencing of only the putative causal gene in a large number of individuals. In one such study, the P-value resulting from the initial exome sequencing is significant enough to suggest causality, and the follow-up sequencing essentially serves to establish the proportion of cases in which the phenotype is attributed to variants in the gene [55]. In others studies, however, the initial sequencing merely identifies potential candidate genes, and follow-up sequencing is required to achieve genome-wide significance [53, 56, 70–72]. In these cases, the second round of sequencing is not corrected for multiple testing, because only a single gene is interpreted during follow-up sequencing. (Additional file 7) Statistical significance of variants found to be causal in selected previous studies.

The rankings of frequencies at which a gene contains rare, deleterious variants is comparable to previously published gene ranking methods for prioritizing variants. The list of genes sorted and ranked according to the number of individuals carrying rare (MAF < 0.5%) heterozygous, loss-of-function variants correlates well with genes ranked based on Z scores calculated from the ExAC dataset consisting of exome sequencing data from 60,706 individuals (ρ = 0.485) [8, 73]. Z scores describe the deviation of observed variant counts per gene from expectation, indicating transcripts that are more intolerant of variation. Despite the fact that 48.8% of genes contain no data (i.e., they contain no variants under these filtering thresholds), ExAC rankings correlate more closely with SORVA rankings than rankings based on RVIS [5] (ρ = 0.374) and FLAGS [7] (ρ = 0.279) methods.

## Discussion

We demonstrate the utility of using mutational burden data to aid in prioritizing variants *in silico* and quantifying the significance of seeing a variant within a gene. We have shown this using examples from previous studies encompassing multiple NGS study designs and disease inheritance models.

Although there was some variation between the frequency of individuals with a rare variant in a given gene between populations, and selecting a comparable population to a study would be ideal when calculating variant significance, this restriction is not necessary. To illustrate, if individuals in the African population frequently carry LOF variants in a gene but this does not hold true for another population that more closely matches the study population, one may nevertheless consider the gene to be less likely to cause a rare Mendelian disorder.

A limitation of this method of ranking genes is that genes are prioritized on the basis of their likelihood of being involved in disease in general rather than in the specific disease of interest [4]. On the other hand, this can be viewed as a benefit in the sense that results are unbiased and do not depend on previously existing annotations, which would bias rankings to prefer known and well-studied genes. This bias is a known issue in the interpretation of clinical variants [74]. To illustrate, Bell et al. discovered that an unexpected proportion (27%) of literature-annotated disease variants in recessive disease-causing genes were incorrect [75], and Piton et al. estimated that 25% of X-linked intellectual disability genes are incorrect or require further review based on allele frequency estimates that have become more accurate with the availability of large-scale sequencing datasets [76]. Disease genes that are incorrectly annotated as disease-causing may explain the lack of difference between the average number of individuals carrying variants in genes causal for autosomal dominant and autosomal recessive genes. One would expect decreased counts for autosomal dominant disease genes due to stronger purifying selection among deleterious variants that arise in these genes, where a single variant may be sufficient to cause disease [77]. Another possibility is that the sample size may be too small to include a sufficient number of individuals who are carriers for rare, deleterious variants in recessive disease genes.

Future improvements to our methods would include increasing the amount of genetic information from unaffected individuals. Our results suggest that for most applications, low MAF thresholds should be used to achieve power to detect genes associated with disease; however, at thresholds of MAF < 0.0005, most genes will lack any data; e.g. there will be no individuals observed who are carriers of LOF variants. The SORVA dataset is useful in its current state with data from a relatively small number of individuals, but increasing the population size by several orders of magnitude will increase the utility of the application. The recently approved Precision Medicine Initiative will fund sequencing and data collection from 1 million or more Americans and make the data accessible to qualified researchers, and the methods described in this manuscript could be applied to this larger dataset and contribute towards the aim of this initiative to generate knowledge applicable to the whole range of health and disease [78].

Additional improvements would include incorporating additional information regarding specific categories of variants, such as the degree to which stop codon gain (also know as nonsense) variants in a gene are constrained to the end of the gene. Knowing whether an essential gene is highly intolerant of nonsense mutations in only certain regions of the gene would allow one to lower the priority of nonsense variants in mutationally tolerant regions when evaluating variants *in silico*. For example, Li et al. exclude stop-gain variants occurring in the terminal gene exon and those that do not affect all transcripts of a gene when evaluating deleterious LOF mutations in a large cohort of individuals [79]. The limitation to providing individual-level mutational burden counts at such a high level of granularity is that researchers will be restricted to following the same methods of filtering and annotating variants. This would be problematic because, by default, many commonly-used software pipelines do not annotate variants with the information about the proportion of transcript truncated [80,–85]. Selecting variant filtering thresholds in SORVA that are identical to those used in one's study is essential in having comparable data with which to calculate variant significance. For this reason, we also did not filter missense variants based on annotations from commonly tools such as SIFT [86], PolyPhen-2 [87], and CADD [88], which provide an interpretation of mutation impacts.

## Conclusions

Our methods provide a score for prioritizing variants within a gene that is unbiased and directly interpretable. Restricted by the sample size of our dataset, we provide limited population-level data, and adding more data will greatly improve the utility of our method. However, even in its current state, SORVA is useful for vetting candidate genes from NGS studies and allows researchers to calculate the significance of seeing a variant in a given gene or protein domain, which is an important step towards developing a quantitative, statistics-based approach for presenting clinical findings.

## Methods

Given the number of technical controls who have a potentially damaging variant anywhere in a gene, we calculated the significance (α < 0.05) of seeing the observed number of singletons (variants observed in a single individual), doubletons (variants observed in two individuals within a single family) and/or tripletons in cases. Specifically, let *p* be the a priori probability that an individual has a heterozygous mutation in a gene. Then, the probability of seeing *N* or fewer singleton/doubleton/tripleton (*i* = 1, 2 or 3, respectively) families with a heterozygous variant, out of *n* total families of identical relatedness, where *r* is the fraction of the genome shared between affected family members is

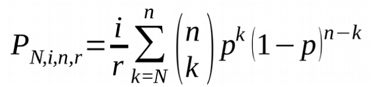

and the significance of seeing rare damaging mutations in, for example, *x* number of singletons and *y* number of doubletons where individuals share ¼ of their genome is calculated as

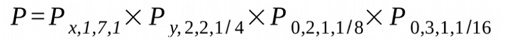

given the family structures of the individuals we had sequenced.

### Datasets

Genomic data and allele frequencies for calculating a priori probabilities of observing a variant within a gene were obtained from the 1000 Genomes Project (phase 3 variant set) [11]. This variant set contains 2504 individuals from 26 populations in Africa (AFR), East Asia (EAS), Europe (EUR), South Asia (SAS), and the Americas (AMR).

### Bioinformatics pipeline

Genomic annotations were assigned to each variation using *SNP & Variation Suite (SVS) v8.1 [80]* with the following parameters: gene set Ensembl release 75 [89], human genome version GRCh37.p13. Variants were filtered for coding mutations that result in a change in the amino acid sequence (e.g. missense, nonsense and frameshift mutations), or mutations that reside within a splice site junction (intronic distance of 2 base pairs). Biallelic data was recoded based on an additive model to correct for MAF of variants on the X chromosome for male samples, using a script in SVS. Variants were then filtered for minor allele frequency thresholds of MAF < 5%, < 1%, < 0.5%, < 0.1% and < 0.05%, based on allelic frequency within the dataset. For each filtered list of variants, we collapsed variants by gene and performed the following two scenarios: 1) an individual was counted as having a rare variant in a gene if the variant mapped to any transcript of a gene; 2) we counted the number of variants in a given gene per individual, i.e. if an individual carried two rare mutations within a gene, they were counted twice. In a separate analysis, we collapsed variants by protein domains obtained from Interpro [13] using the Ensembl API [82]. Finally, we repeated each analysis using a subset of the 1000 Genomes Project data grouped according to superpopulation. Variant collapsing methods were performed using a custom Python script run by SVS, and an individual was counted as having a rare variant in a gene if the variant mapped to any transcript of a gene.

In addition to replicating the analysis for gene versus protein domain, for each population, and for each MAF threshold, we also repeated the calculations for multiple categories of predicted variant consequence on the protein transcript. The two categories were 1) nonsynonymous variants or those predicted to be more severe by Ensembl [89], briefly nonsynonymous or LOF variants, and 2) potential LOF variants (includes splice site, protein truncation stop codon gain mutations, and frameshift indels).

### Comparison of gene ranking methods

Genic mutational intolerance scores were obtained from three previous studies and included the Residual Variation Intolerance Score (RVIS) [5], scores from Shyr et al. 2014 (FLAGS) [7] and Z scores based on the ExAC dataset [73, 90]. We considered 16,302 genes that were found in all three datasets, as well as ours, and ranked genes based on scores obtained using each method. Spearman's rho test [91, 92] was used to measure the size and statistical significance of the association between the rankings obtained from ExAC and those obtained by RVIS, FLAGS and SORVA methods. This test measures the strength and direction of association between two ranked variables.

### Comparison of disease gene categories

To determine whether our results show concordance with studies identifying essential genes critical for the survival of a human, we compared the number of individuals with rare, deleterious mutations between gene lists containing essential human genes, those known to cause Mendelian diseases, and control genes, defined as genes not included in either category. We considered genes to be essential human genes if they were determined as such in at least one of the following two studies. The first essential human gene set is defined as 'core' essential genes that are required for fitness of cells from both the HAP1 and KBM7 cell lines, determined through extensive mutagenesis in near-haploid human cells (N=1734) [93]. The second essential human gene set consists of genes essential to four screened cell lines, KBM7, K562, Raji and Jiyoye, determined using the CRISPR system. From the latter set, we selected genes with an adjusted P-value CRISPR score< 0.4025 for each cell line (N=1878) [94].

To identify genes known to cause Mendelian disease, we parsed data from Online Mendelian Inheritance in Man (OMIM) [1] and identified phenotype descriptions with known molecular basis. We parsed the genotype description field for the gene name and the following phrases: ' caused by heterozygous/homozygous mutation', 'autosomal recessive', 'autosomal dominant', 'X-linked', ' on chromosome X', and categorized genes as autosomal recessive (AR) (N=655), autosomal dominant (AD) (N=785), and X-linked (XL) (N=126) accordingly.

### Calculating depletion of variants in protein domains

We performed two analyses: first, we calculated whether protein domains in a gene were depleted of variation compared to the rest of the gene, and second, we calculated whether there were any types of protein domains that were depleted of variation in general across the entire genome.

First, for each protein domain mapping within a gene, we calculated whether domains were depleted of variation compared to the rest of the gene. Depletion was calculated as: (number of variants per individual in protein domain / number of variants per individual in gene × length of protein domain / length of transcript). A value of 1 is expected by chance, and a small value indicates protein domains most intolerant towards mutations. We then calculated the *P*-value of obtaining such a depletion score using the binomial cumulative density function, under the assumption that each site is equally likely to be mutated. This *P*-value is then "PHRED-scaled" by expressing the rank in order of magnitude terms rather than the precise rank itself. High scaled scores indicate that a protein domain is depleted of rare (MAF < 0.5%) mutations compared to the rest of the gene, hence protein domains with high scores tend to be enriched for highly mutated genes. We filtered out genes with no observed mutations and protein domains that span more than 50% of the length of the transcript, resulting in 7,828 genes remaining.

Next, we calculated whether there were any types of protein domains that were depleted of variation in general across the entire genome. We weighted each gene with instances of the protein domain equally. In other words, if a gene had multiple instances of a protein domain, we first calculated the mean number of heterozygous rare (MAF <=0.5%) LOF variants observed (in the entire dataset of 2,504 individuals) in either protein domain within the gene. Next, we calculated the mean and variance of the means for each gene.

To determine whether a protein domain was well covered by sequencing, we calculated the mean coverage of an instance of a protein domain in the 1000 Genomes Project sample HG00096. We calculated depth of coverage from phase 3 exome alignment data using GATK and custom code, which is available at https://github.com/alizrrao/DepthOfCoveragePerInterval.

## Declarations

### Author’s contributions

ARR and SFN designed analyses. ARR analyzed data and created the software. ARR and SFN wrote the manuscript.

### Author details

^1^Department of Human Genetics, University of California, Los Angeles, Los Angeles, California, USA.

^2^Department of Psychiatry and Biobehavioral Sciences at the David Geffen School of Medicine, University of California, Los Angeles, Los Angeles, California, USA. ^3^Department of Pathology and Laboratory Medicine, University of California, Los Angeles, Los Angeles, California, USA.

### Availability of data and materials

Gene-based mutational burden datasets and the webtool are available for querying at the SORVA website, https://sorva.genome.ucla.edu.

Standalone software and datasets are freely available for download at https://github.com/alizrrao/sorva.

The 1000 Genomes Project datasets analysed during the current study are available in the International Genome Sample Resource (IGSR), http://www.internationalgenome.org/data.

## Acknowledgements

Funding to ARR was provided by National Institutes of Health (NIH) Training Grant No. T32HG002536. We thank the NIH/National Center for Advancing Translational Science (NCATS) (UCLA CTSI Grant Number UL1TR000124) for their support. The funders had no role in study design, data collection and analysis, decision to publish, or preparation of the manuscript.

## Additional Files

**Table 2. Statistical significance of variants found to be causal in selected previous studies.** Applying our methods to previous NGS findings, in which researchers filtered variants using various criteria, would have statistically validated findings *in silico*. See Additional file 7 for more details. ^†^The parameter *f* denotes the proportion of individuals in the 1000 Genomes Project dataset who have a rare variant at least as severe as the identified variants. ^&^We used a threshold of MAF < 0.1% for studies with no specific MAF threshold. A MAF threshold labeled exclusion refers to studies where variants were not filtered for a given threshold and variants were excluded based on their presence in public databases such as dbSNP. *Follo-up Sanger sequencing identified mutations in 2 out of 3 exome-negative cases. Follow-up sequencing of the given gene identified further mutations in multiple additional cases. Abbreviations: MAF, minor allele frequency; AD, autosomal dominant; AR, autosomal recessive; XL, X-linked; nonsyn, nonsynonymous variant; LOF, loss-of-function variant; Het, heterozygous; Hom, homozygous; CHet/Hom, compound heterozygous or homozygous.

**Additional file 1: Number of individuals carrying a rare variant in a gene under various filtering thresholds.** Each data point represents a single gene which contains a variant in the aggregate population (n=2504 individuals). Calculations were repeated using multiple variant filtering thresholds to determine the scenario that most differentiates between essential genes, those known to cause autosomal dominant, autosomal recessive or X-linked disease, and other genes. We varied filters for type of variant (‘LOF or missense’ or ‘LOF only’), zygosity (Het or Hom) and MAF threshold. Colored shapes indicate the centroids of each group of genes. Abbreviations: LOF, loss-of-function; nonsyn, nonsynonymous or LOF; het, heterozygous; hom, homozygous; ess, essential; AD, autosomal dominant; AR, autosomal recessive; XL, X-linked.

**Additional file 2: Mean number of individuals mutated for different types of protein domains.** We calculated the mean number of individuals (out of 2504 individuals) who carried mutations in a given type of protein domains, averaging per gene.

**Additional file 3: Variant depletion scores for all protein domain in every gene.** For each instance of a protein domain in a gene, we calculated variant depletion scores to identify regions within a gene that may be under differing degrees of evolutionary constraint.

**Additional file 4: Supplementary methods.** Includes derivation of equations and math used for calculating the significance of finding rare variants in a given gene.

**Additional file 5: Screenshot of an example query run on SORVA.** Users can select variant filtering thresholds such as population, MAF cutoff, zygosity and whether to consider only LOF variants or missense variants, as well. Output includes the number of individuals who carry a rare variant in the gene and in any protein domain that maps to the gene.

**Additional file 6: List of candidate autism genes.** Genes listed were used to produce Figure 5.

**Additional file 7: Calculating P-values for findings from previous whole-exome or targeted sequencing studies.** The parameter *f* denotes the proportion of individuals in the 1000 Genomes Project dataset who have a rare variant at least as severe as the identified variants. A MAF threshold labeled exclusion refers to studies that did not filter by a given threshold and excluded variants based on their presence in public databases such as dbSNP; in such cases, results were calculated using a MAF threshold of 0.1%. Abbreviations: MAF, minor allele frequency; AD, autosomal dominant; AR, autosomal recessive; XL, X-linked; nonsyn, nonsynonymous variant; LOF, loss-of-function variant; Het, heterozygous; Hom, homozygous; CHet/Hom, compound heterozygous or homozygous.

